# The S1 protein of SARS-CoV-2 crosses the blood-brain barrier: Kinetics, distribution, mechanisms, and influence of ApoE genotype, sex, and inflammation

**DOI:** 10.1101/2020.07.15.205229

**Authors:** Elizabeth M. Rhea, Aric F. Logsdon, Kim M. Hansen, Lindsey Williams, May Reed, Kristen Baumann, Sarah Holden, Jacob Raber, William A. Banks, Michelle A. Erickson

## Abstract

Evidence strongly suggests that SARS-CoV-2, the cause of COVID-19, can enter the brain. SARS-CoV-2 enters cells via the S1 subunit of its spike protein, and S1 can be used as a proxy for the uptake patterns and mechanisms used by the whole virus; unlike studies based on productive infection, viral proteins can be used to precisely determine pharmacokinetics and biodistribution. Here, we found that radioiodinated S1 (I-S1) readily crossed the murine blood-brain barrier (BBB). I-S1 from two commercial sources crossed the BBB with unidirectional influx constants of 0.287 ± 0.024 μL/g-min and 0.294 ± 0.032 μL/g-min and was also taken up by lung, spleen, kidney, and liver. I-S1 was uniformly taken up by all regions of the brain and inflammation induced by lipopolysaccharide reduced uptake in the hippocampus and olfactory bulb. I-S1 crossed the BBB completely to enter the parenchymal brain space, with smaller amounts retained by brain endothelial cells and the luminal surface. Studies on the mechanisms of transport indicated that I-S1 crosses the BBB by the mechanism of adsorptive transcytosis and that the murine ACE2 receptor is involved in brain and lung uptake, but not that by kidney, liver, or spleen. I-S1 entered brain after intranasal administration at about 1/10^th^ the amount found after intravenous administration and about 0.66% of the intranasal dose entered blood. ApoE isoform or sex did not affect whole brain uptake, but had variable effects on olfactory bulb, liver, spleen, and kidney uptakes. In summary, I-S1 readily crosses the murine BBB, entering all brain regions and the peripheral tissues studied, likely by the mechanism of adsorptive transcytosis.

**Graphical Abstract:** 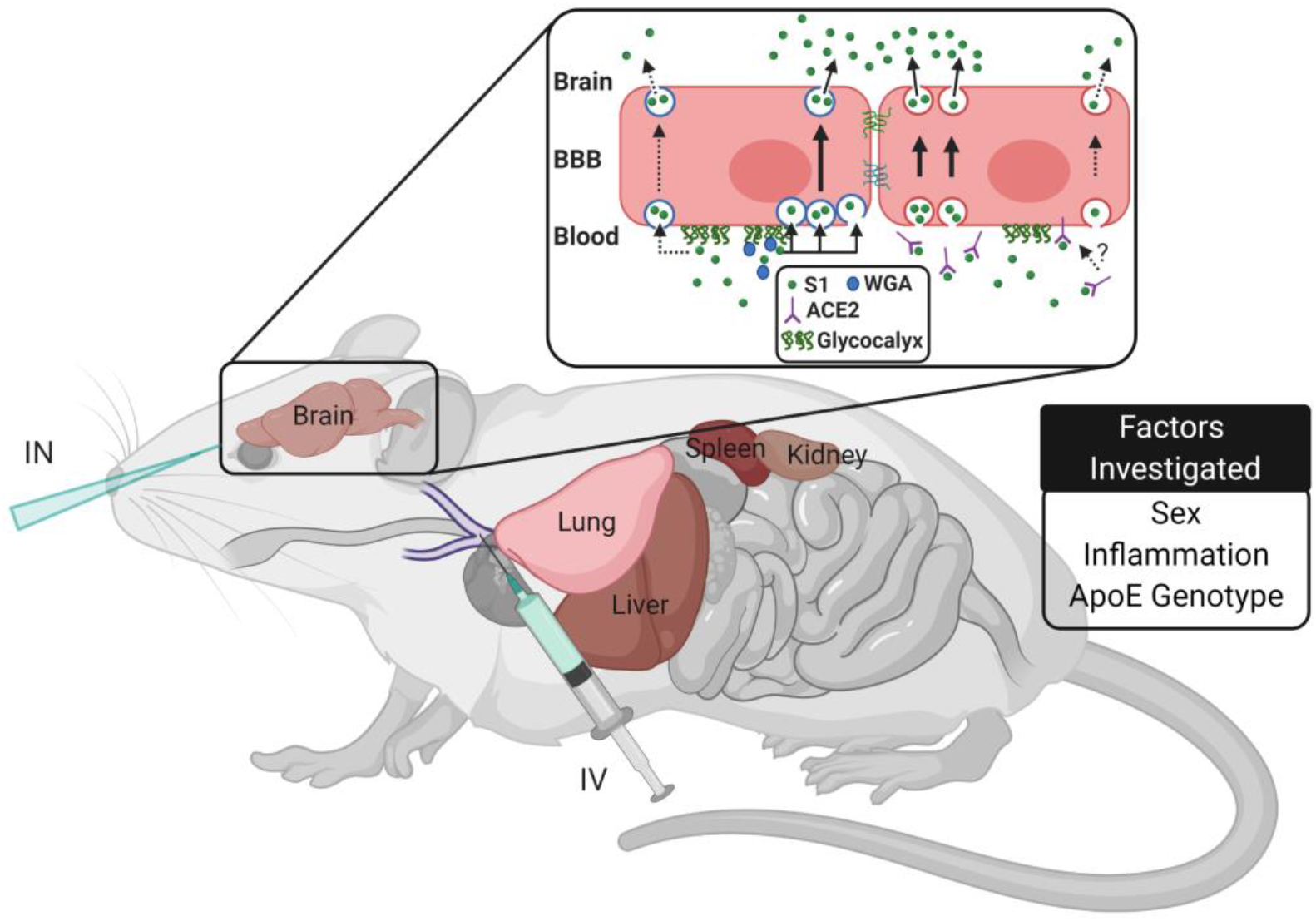

## Introduction

Severe acute respiratory syndrome corona virus 2 (SARS-CoV-2) is responsible for the COVID-19 pandemic. In addition to pneumonia and acute respiratory distress, COVID-19 is associated with a host of symptoms that relate to the central nervous system (CNS), including loss of taste and smell, headaches, twitching, seizures, confusion, vision impairment, nerve pain, dizziness, impaired consciousness, nausea and vomiting, hemiplegia, ataxia, stroke, and cerebral hemorrhage ^1, 2^. It has been postulated that some of the symptoms of COVID-19 may be due to direct actions on the CNS, such as impaired respiratory symptoms being in part due to SARS-CoV-2 invading the respiratory centers of the brain ^1, 3^. SARS-CoV-2 mRNA has been recovered from the cerebrospinal fluid ^4^, strongly suggesting that SARS-CoV-2 can cross the blood-brain barrier (BBB). Other evidence also suggests that SARS-CoV-2 is to cross the BBB. Encephalitis has been reported, which could be a result of virus, viral proteins, or severe systemic inflammation ^4, 5^. Other corona viruses, including the closely related SARS virus that caused the 2003-4 outbreak, are able to cross the BBB ^6–8^ and SARS-CoV-2 can infect neurons in a brainsphere model ^9^.

CNS symptoms do not prove viral entry. There are many mechanisms by which viruses can induce changes in the CNS without themselves directly crossing the BBB. First, COVID-19 is associated with a cytokine storm and many cytokines cross the BBB to affect CNS function ^10^. Secondly, viral proteins as exemplified by gp120 and the tat protein of HIV-1 are often circulating in blood free of virus or cells and, having the ability to themselves cross the BBB, can produce neuroinflammation, activate the hypothalamic-pituitary-adrenal axis, and otherwise harm CNS functions ^11–17^.

Thus, the study of the ability of SARS-CoV-2 viral proteins to cross the BBB is important and clinically relevant for several reasons. First, the ability of those proteins to reach the CNS can itself be a source of CNS symptoms and events. Second, study of the viral proteins can be predictive of the rate, distribution, and mechanisms of passage across the BBB of the whole virion ^18, 19^. Third, use of the protein of a virus allows study of biodistribution in the absence of replicating viral infection and to readily assess the effects of risk factors such as sex and ApoE status on rates of uptake and biodistribution ^20, 21^. Replicating viral infection often results in secondary phenomena such sickness behavior, widespread organ failure, and inflammation, making it difficult to separate actions of the virus from the reactions to the virus. Fourth, the species barrier is often not a major factor when one can directly study the pharmacokinetics of virus or viral proteins using methods that do not depend on viral infection. For example, the species barrier for HIV-1 exists because the mouse lacks critical elements needed for HIV-1 replication ^22, 23^, but HIV-1 does penetrate the mouse BBB ^24^. Fifth, it is very difficult to demonstrate the pathway taken from a site of inoculation to a site of invasion when viral infection is used. For example, virus placed within the nasal cavity could reach the brain by way of retrograde entry into the CNS by way of the olfactory bulb, the trigeminal nerves, or by hematogenous spread. With labeled virus or viral proteins, these pathways can be precisely compared for the initial inoculant.

Coronavirus spike proteins are the major determinants of their tropism and pathogenicity ^25^. The S1 subunit of spike contains a receptor binding domain that binds to cell surface receptors and facilitates subsequent viral fusion with the cell membrane. Recently, it has been shown that the SARS-CoV-2 spike protein binds angiotensin converting enzyme 2 (ACE2) which mediates entry of the virus into cells ^26–28^. Here, we found that the radioiodinated S1 protein from SARS-CoV-2 (I-S1) from two different commercial sources readily crossed the murine BBB. We found it entered the parenchymal tissue of brain and to a lesser degree was sequestered by brain endothelial cells and associated with the brain capillary glycocalyx. We measured its rate of entry into brain after intravenous and intranasal administration, determined uptake by 11 different brain regions, examined the effect of inflammation, ApoE status, and sex on transport and compared brain uptake to the uptakes by liver, kidney, spleen, and lung. Based on studies with the glycoprotein wheatgerm agglutinin, we found brain entry of I-S1 was likely involved the vesicular-dependent mechanism of adsorptive transcytosis.

## Methods

### Mice

All mouse studies were approved by the local IACUC at OHSU and the VA Puget Sound Healthcare System and conducted in AAALAC approved facilities. Male CD-1 mice (6-10 weeks old) were purchased from Charles River Laboratories (Hollister, CA). Human E3- and E4-targeted replacement (TR) male and female mice, generated as described ^29^, were bred at the Oregon Health & Sciences University (OHSU) prior to transfer to the Veterans Affairs Puget Sound Health Care System (VAPSHCS) for the experiments of this study. E3 and E4 TR mice were approximately 4 months of age on the day of the study. Mice had *ad libitum* access to food and water and were kept on a 12/12 hour light/dark cycle. For all studies, mice were anesthetized with an intraperitoneal (ip) injection of 0.15-0.2 mL of 40% urethane to minimize pain and distress. Anesthetized mice were placed on a heating pad until time of use. At the end of each study, mice were euthanized by decapitation while under anesthesia.

### Radioactive Labeling of Proteins

The S1 proteins (Raybiotech, Cat no 230-30161, Peachtree Corners, GA; Amsbio, AMS.S1N-C52H3, Abingdon, UK) were provided by the manufacturer dissolved in phosphate buffered saline (PBS, pH 7.4) at a concentration ranging from 0.45-0.61 mg/mL. Upon receipt, the S1 proteins were thawed and aliquoted into 5μg portions and either used immediately or stored at −80 °C until use. The 5 μg of thawed S1 protein was radioactively labeled with 1 mCi 125I (Perkin Elmer, Waltman, MA) using the chloramine-T method ^30^, as described and the radioiodinated S1 (I-S1) purified on a column of G10 Sephadex (GE Healthcare, Uppsala, SE), eluting with phosphate buffer (PB) into glass tubes containing 1% bovine serum albumin in lactated Ringer’s solution (BSA-LR). Bovine serum albumin (Sigma, St. Louis, MO) was labeled with ^99m^Tc (GE Healthcare, Seattle, WA) using the stannous tartrate method and the ^99m^Tc – labeled albumin (T-Alb) purified on a column of G-10 Sephadex. Both of the I-S1’s and the T-Alb were more than 90% acid precipitable. The molecular weight of the labeled proteins was further confirmed by running 200,000-600,000 CPM activity in 1x LDS buffer (Invitrogen) with or without reducing agent (Invitrogen) on a 4-12% bis-tris gel (Genescript) in 3-(N-morpholino)propanesulfonic acid (MOPS) buffer (Invitrogen). The gel was then fixed for 30 min in 10% acetic acid/50% methanol, washed 3x with water, and then dried using a DryEase^®^ Mini-Gel Drying System (Invitrogen). Dried gels were exposed on autoradiography film for 24 hours and then developed. Supplemental Figure 1 shows that major bands of I-S1 from Raybiotech and Amsbio migrated at their predicted molecular weight patterns, based on manufacturer data.

### Multiple-time Regression Analysis (MTRA)

In anesthetized mice, the left jugular vein was exposed for an intravenous (iv) injection of 0.1 mL BSA-LR containing 3×10^5^ cpm of I-S1 and 6×10^5^ cpm of T-Alb. At time points between 1 and 30 min, blood was collected from the carotid artery. Blood was centrifuged at 3200xg for 10min and 50uL serum was collected. The whole brain, kidney, and spleen and portions of the lung and liver were removed and weighed. Tissues and serum were placed into a Wizard 2 gamma counter (Perkin Elmer), and the levels of radioactivity were measured. Results for serum were expressed as the percent of the injected dose per mL of blood (%Inj/mL). Results for brain and tissues were expressed as the tissue/serum ratio in units of μL/g. For each individual tissue, its ratio for T-Alb was subtracted from its ratio for I-S1, yielding the “delta” value which reflects values corrected for vascular space and any nonspecific leakage into tissue. The delta brain/serum ratios were plotted against exposure time (Expt), a calculation that corrects for clearance from blood:

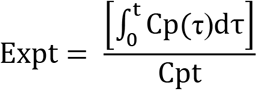

where t is the time between the iv injection and sampling, Cp is the cpm/mL of arterial serum, Cpt is the cpm/mL of arterial serum at time t, and *τ* is the dummy variable for time. The slope of the linear portion of the relation of tissue/serum ratio vs Expt measures the unidirectional influx rate (Ki in units of μL/g-min) and the Y-intercept measures Vv, the vascular space^31, 32^.

The area under the curve for the level of radioactivity in blood from 0-30 min was calculated using Prism 8.0 software (GraphPad Inc, San Diego, CA). The percent of the injected dose per gram of brain (%Inj/g) was calculated by multiplying the delta brain/serum value by the %Inj/mL for I-S1.

### Stability in Brain and Blood

Anesthetized mice received an injection into the jugular vein of 0.1 mL of BSA-LR containing 6×10^5^ cpm of I-S1 plus 6×10^5^ cpm of T-Alb. Ten minutes after the injection, arterial blood was collected from the abdominal aorta, the thorax opened and the descending thoracic artery clamped, both jugular veins severed, and 20 mL of lactated Ringer’s solution perfused through the left ventricle of the heart to wash out the vascular space of the brain. The whole brain was then removed. Whole blood was centrifuged and 10 uL of serum was added to 500 uL BSA-LR and combined with an equal part of 30% tricholoroacetic acid, mixed, centrifuged, and the supernatant and pellet counted. Whole brain was homogenized in BSA-LR plus complete mini protease inhibitor (Roche, Mannheim, DE; one tab per 10 mL buffer) with a hand-held glass homogenizer, and centrifuged. The resulting supernatant was combined with equal parts of 30% trichloroacetic acid, mixed, centrifuged, and the supernatant and pellet counted. To determine the amount of degradation that occurred during processing, I-S1 and T-Alb was added to brains and arterial whole blood from animals that had not been injected with radioactivity and processed immediately as above. The percent of radioactivity that was precipitated by acid (%P) in all of these samples was calculated by the equation:

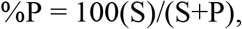

where S is the cpm in the supernatant and P is the cpm in the pellet.

### Capillary Depletion

The capillary depletion method as adapted to mice was used to separate cerebral capillaries and vascular components from brain parenchyma ^33, 34^. We used the variant of the technique that also estimates reversible binding to the capillary lumen. Mice were anesthetized and received an iv injection of 6×10^5^ cpm Tc-Alb with 6×10^5^ cpm I-S1 in 0.1 mL BSA-LR. 5, 10, and 30 min later, blood was obtained from the carotid artery, and the brain (non-washout) was extracted. In other mice, blood was taken from the abdominal aorta at 5, 10, and 30 min, the thorax was opened and the descending thoracic artery clamped, both jugulars severed, and 20 mL of lactated Ringer’s solution infused via the left ventricle of the heart in order to washout the vascular contents of the brain and to remove any material reversibly associated with the capillary lumen. Each whole brain was homogenized in glass with physiological buffer (10mM HEPES buffer, 141 mM NaCl, 4mM KCl, 2.8 mM CaCl_2_, 1 mM MgSO_4_, 1 mM NaH_2_PO_4_-H_2_O, 10 mM D-glucose, pH 7.4) and mixed thoroughly with 26% dextran. The homogenate was centrifuged at 4255xg for 15 min at 4°C. The pellet, containing the capillaries, and the supernatant, representing the brain parenchymal space, were carefully separated. Radioactivity levels in the capillary pellet, the brain supernatant, and the arterial serum were determined for both T-Alb and I-S1 and expressed as the capillary/serum and brain parenchyma/serum ratios. The I-S1 parenchymal ratios were corrected for vascular contamination by subtracting the corresponding ratios for T-Alb; these results are reported as the delta brain parenchyma/serum ratios. The amount of S1 in the brain parenchymal space was taken as the delta brain parenchymal space from the washout group (PW), the amount in the capillary as the capillary from the washout group (CW), and the amount of material loosely binding to the luminal surface (Luminal) as:

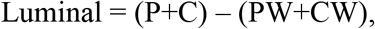

Where P are the delta brain parenchymal space and the capillary space, both from the non-washout groups.

### Effects of Wheatgerm Agglutinin

Mice were anesthetized after which the jugular vein and right carotid artery were exposed. They were then given a jugular vein injection of 0.1 mL BSA-LR containing 3×10^5^ cpm of I-S1 and 6×10^5^ cpm of T-Alb. For some mice, 10 μg/mouse of a plant lectin (wheat germ agglutinin [WGA], Sigma, St. Louis, MO) was included in the iv injection. Brain, tissues, and serum samples were collected 5 min later and tissue/serum ratios calculated.

### Saturation and Modulation of Uptake

In anesthetized mice, the left jugular vein was exposed for an iv injection of 0.1 mL BSA-LR containing 3×10^5^ cpm of I-S1 (Raybiotech) and 6×10^5^ cpm of T-Alb. For some mice, the injection contained 1 μg/mouse of unlabeled S1 (Amsbio), mouse acyl ghrelin (CBio, Menlo Park, CA), angiotensin II (Tocris, Bristol, UK), or human ACE2 (R&D, Minneapolis, MN). Ten min after iv injection, whole blood was obtained from the carotid artery and centrifuged after clotting. The whole brain, olfactory bulb, kidney, spleen, and portions of the liver and lung removed. The levels of radioactivity in the arterial serum and the tissues were determined and the results expressed as the percent of the injected I-S1 per mL for serum and delta tissue/serum ratios (μL/g) for the tissues.

### Effects of Inflammation: Treatment with Lipopolysaccharide

Male CD-1 mice aged 6-10 weeks were given an ip injection of 3 mg/kg LPS from *Salmonella typhimurium* (Sigma, St. Louis, MO) dissolved in sterile normal saline at 0, 6, and 24 h. At 28 h, mice were anesthetized and the left jugular vein and right carotid artery exposed. The mice were given an iv injection of 6×10^5^ cpm of I-S1 and 6×10^5^ cpm T-Alb in 0.1 mL of BSA/LR into the left jugular vein. Arterial blood was collected from the right carotid artery 10 min later, the mouse immediately decapitated, the brain removed, dissected into regions (olfactory bulb, frontal cortex, occipital cortex, parietal cortex, thalamus, hypothalamus, striatum, hippocampus, pons-medulla, cerebellum, midbrain), and the regions weighed. Kidney, spleen, and portions of the liver and lung were also removed and weighed. Serum was obtained by centrifuging the carotid artery blood for 10 min at 4255 xg. Levels of radioactivity in serum, brain regions, and tissues were measured in a gamma counter. Whole brain values were calculated by summing levels of radioactivity and weight for all brain regions except for olfactory bulb. The levels of radioactivity in the arterial serum and the brain regions and tissues were determined and the results expressed as the percent of the injected I-S1 per mL for serum and delta tissue/serum ratios (μL/g) for the tissues.

### Intranasal Delivery

Anesthetized mice were placed in the supine position and received a 1 μL injection of 2×10^5^ - 3×10^5^ cpm I-S1 in BSA/LR administered to each naris, delivered to the level of the cribriform plate (4 mm depth), using a 10 μL Multi-flex tip (Thermo Fisher Scientific, Waltham, MA). After administration, the mouse remained in the supine position for 30 s before being placed on the left side. Arterial blood was collected from the right carotid artery 10 or 30 min later, the mouse immediately decapitated, the brain removed, dissected into regions (olfactory bulb, frontal cortex, occipital cortex, parietal cortex, thalamus, hypothalamus, striatum, hippocampus, pons-medulla, cerebellum, midbrain), and the regions weighed. Serum was obtained by centrifuging the carotid artery blood for 10 min at 4255 xg. Levels of radioactivity in serum and brain regions were measured in a Wizard 2 gamma counter. Whole brain values were calculated by summing levels of radioactivity and weight for all brain regions except for olfactory bulb. The levels of radioactivity in the arterial serum and the tissues were determined and the results expressed as the percent of the injected I-S1 per mL for serum and the percent of the injected I-S1 per g of brain region calculated.

### MTRA in ApoE mice

Multiple-time regression analysis as described above was performed in female and male homozygous human E3- and E4-targeted replacement (TR) mice. Two-way ANOVA used sex and ApoE isoform as independent variables.

### Transport by iPSC-derived brain endothelial-like cells (iBECs)

The iBECs were derived from the GM25256 iPSC line (Coriell Institute) using the method of Neal et al ^35^ with a seeding density of 15,000 cells per well for differentiation which was found to be optimal for this cell line. Briefly, iPSCs were grown to optimal density on plates coated with Matrigel (VWR cat no. 62405-134) in E8 Flex medium (Thermo Fisher Scientific cat no. A2858501), and then passaged using Accutase (Thermo Fisher Scientific cat no. A1110501) onto Matrigel-coated plates in E8 Flex medium plus 10 μM ROCK inhibitor Y-27632 (R&D Systems, cat no. 1254). The next day, the medium was changed to E6 (Thermo Fisher Scientific, cat no. A1516401) and E6 changes continued daily for 3 more days. Next, the medium was changed to human endothelial serum-free medium (HESFM, Thermo Fisher Scientific, cat no. 11111044) supplemented with 20 ng/mL bFGF (Peprotech, cat no. 100-18B), 10 μM retinoic acid (Sigma, cat no. R2625), and 1% B27 supplement (Thermo Fisher Scientific, cat no. 17504044). 48 h later, iBECs were subcultured onto 24-well transwell inserts (Corning cat no. 3470) coated with 1 mg/mL Collagen IV (Sigma, cat no. C5533) and 5mM Fibronectin (Sigma, cat no. F1141) in HESFM + 20 ng/mL bFGF, 10 uM retinoic acid, and 1% B27. 24 h after subculture, the medium was changed to HESFM + 1% B27 without bFGF or retinoic acid, and transendothelial electrical resistance (TEER) was recorded using an End EVOM2 Voltohmmeter (World Precision Instruments, Sarasota Florida) coupled to an ENDOHM cup chamber. TEER measurements occurred daily and S1 transport experiments were conducted when TEER stabilized, between 10-13 days in vitro.

Prior to transport studies, the medium was changed and cells were equilibrated in the incubator for 20 min. Warm HESFM + 1% B27, 1 million CPM T-Alb, and 500,000 CPM of I-S1 was then added in a volume of 100 μL to the luminal chamber. After incubation times of 10, 20, 30, and 45 min at 37 °C, 500 μL volumes of medium from the abluminal chamber were collected and replaced with fresh pre-warmed medium. Samples were then acid precipitated by adding a final concentration of 1% BSA to visualize the pellet and 15% trichloroacetic acid to precipitate the proteins in solution. The samples were centrifuged at 4255xg for 15 min at 4 °C. Radioactivity in the pellet was counted in the gamma counter and the permeability-surface area coefficients for T-Alb and I-S1 were calculated according to the method of Dehouck et al ^36^. Clearance was expressed as microliters (μL) of radioactive tracer that was transported from the luminal to abluminal chamber, and was calculated from the initial level of acid-precipitable radioactivity added to the luminal chamber and the final level of radioactivity in the abluminal chamber:

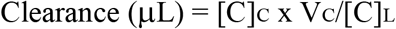

Where [C]_L_ is the initial concentration of radioactivity in the luminal chamber (in units of CPM/μL), [C]_C_ is the concentration of radioactivity in the abluminal chamber (in units of CPM/μL) and Vc is the volume of the abluminal chamber in μL. The volume cleared was plotted vs. time, and the slope was estimated by linear regression. The slopes of clearance curves for the iBEC monolayer plus Transwell^®^ membrane was denoted by PS_app_, where PS is the permeability × surface area product (in μL/min). The slope of the clearance curve for a Transwell^®^ membrane without iBECs was denoted by PS_membrane_. The PS value for the iBEC monolayer (PS_e_) was calculated from 1 / PS_app_ = 1 / PS_membrane_ + 1 / PS_e_. The PS_e_ values were divided by the surface area of the Transwell^®^ inserts (0.33 cm^2^) to generate the endothelial permeability coefficient (Pe, in μL/min/cm^2^).

### Statistics

Means are reported with their SE and n. Two-tailed t-tests were used to compare two means and analysis of variance (ANOVA) followed by multiple comparisons test when more than two means are compared. The Prism 8.0 statistical software package was used for all statistical calculations (GraphPad Inc, San Diego, CA). Linear regression lines were calculated and their slopes and intercepts statistically compared using the program in Prism 8.0 software. As required by multiple-time regression analysis, only the linear portion of the slope was used to calculate Ki. Outliers whose exclusion improved the r^2^ > 0.2 were excluded from analysis. More than two slopes were compared by ANOVA; the standard error of the slope computed by the Prism program was used and the degrees of freedom being n - 2.

## Results

Figure 1 compares the I-S1 proteins from two sources, Raybiotech and Amsbio, for their abilities to cross the BBB as assessed by MTRA. Also shown are values for the vascular space marker T-Alb which acted as an internal control for each animal studied. For this figure only, to facilitate comparison to T-Alb, brain/serum ratios rather than delta brain/serum ratios are shown. There was a strong, positive relation between brain/serum ratios and Expt for both I-S1 proteins: r^2^ = 0.935, p<0.0001 (Raybiotech); r^2^ = 0.905, p<0.0001 (Amsbio) demonstrating transport across the BBB. The unidirectional influx rates (Ki’s) for both compounds were nearly identical: 0.295 ± 0.022 μL/g-min (Raybiotech) vs 0.304 ± 0.033 μL/g-min (Amsbio). The Vi values were also similar: 8.74 ± 0.50 μL/g (Raybiotech) vs 6.88 ± 0.84 μL/g (Amsbio). Comparison of brain/serum ratio vs Expt lines revealed no statistical difference between the Ki’s, but there was a statistical difference in the Vi’s: F(1,23) = 10.32, p = 0.0039, with the Vi for Ambsio I-S1 being lower. T-Alb showed no correlation between its brain/serum ratios and Expt and so its values only reflected the vascular space of the brain.

**Figure 1.**
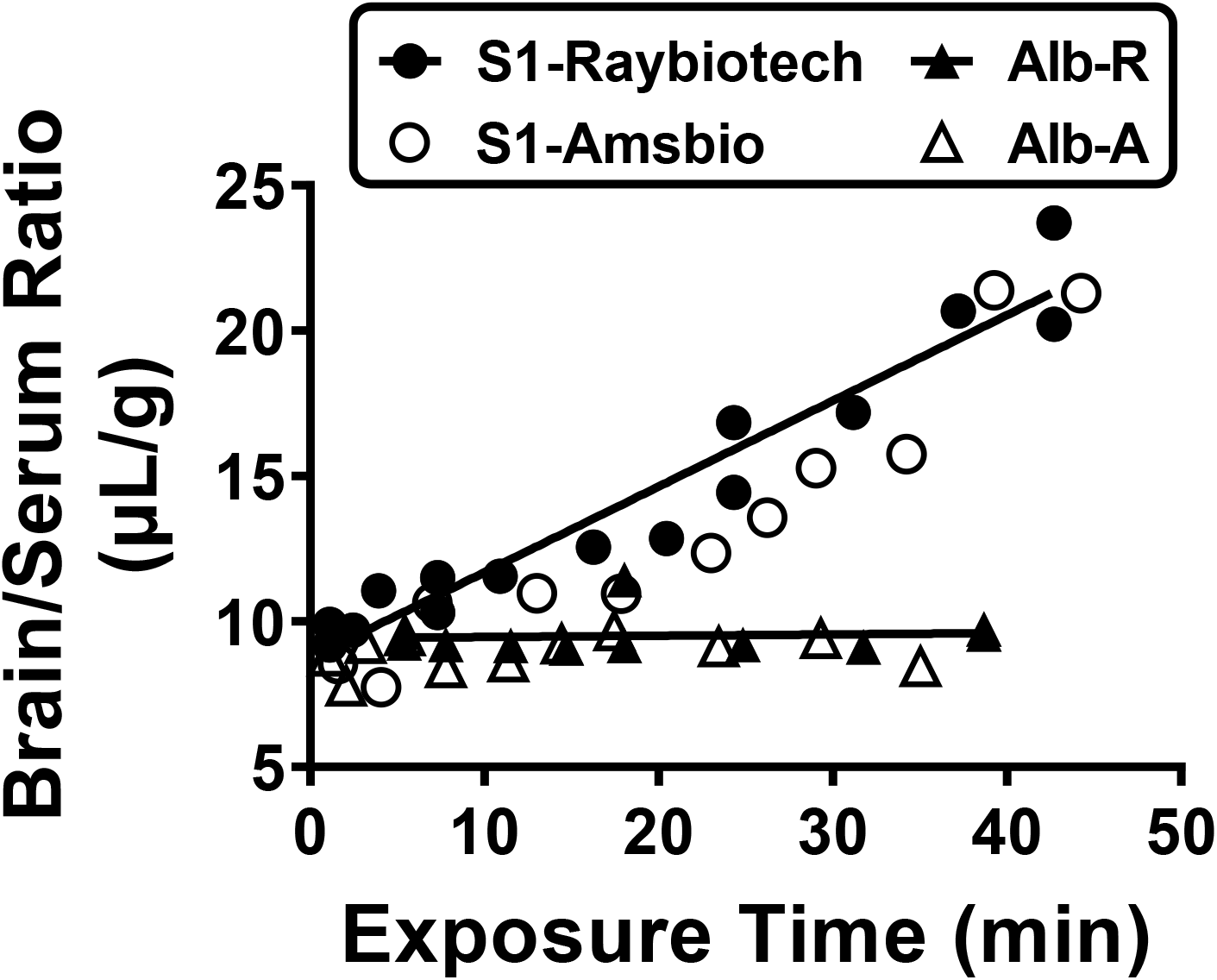
Comparison of BBB transport of S1 proteins from two sources, Raybiotech and Amsbio. Co-injected T-Alb (triangles) measured vascular space of brain and leakage and showed no evidence of BBB transport. Both S1 proteins (circles) crossed the BBB with no differences in the transport rates (Ki), but with a lower Vi for Amsbio S1. See Results for statistical and pharmacokinetic parameters.

Figure 2 shows the clearance of I-S1 (Raybiotech) from blood and its uptake by brain and other tissues. Clearance from blood was linear for the first 10 min with a half-time clearance of about 6.6 min (Figure 2A). For the entire 30 min of the experiment, data was fit to a one phase decay model with Y0 = 1.75, plateau = 1.13, K = 0.165, and half life = 4.20 min, r^2^ = 0.956. The delta tissue/serum ratios vs Expt for brain and other tissues; that is, tissue/serum ratios that have been corrected for T-Alb, are shown in Figure 2B-F. T-Alb measures both the vascular space of the tissue as well as any leakage into the tissue bed. Since I-S1 is about twice the size of T-Alb and leakage is usually a function of the inverse square root of the molecular weight, T-Alb is an appropriate control for leakage of I-S1 as well. Delta tissue/serum ratios, therefore, represent the selective uptake of I-S1. Brain had a Ki = 0.287 ± 0.024 μL/g-min, r^2^ = 0.938, p<0.0001 (Figure 2B). Uptake patterns and rates are shown in figure 2 for lung (2C, r^2^ = 0.903, p<0.0001), spleen (2D, r^2^ = 0.838, p = 0.0014), kidney (2E, r^2^ = 0.335, p = 0.024), and liver (2F, r^2^ = 0.816, p = 0.0001). Brain, lung, and kidney showed linear uptake throughout the study. Spleen and liver had nonlinear relations with time, consistent with an efflux of S1 back into the blood stream.

**Figure 2.**
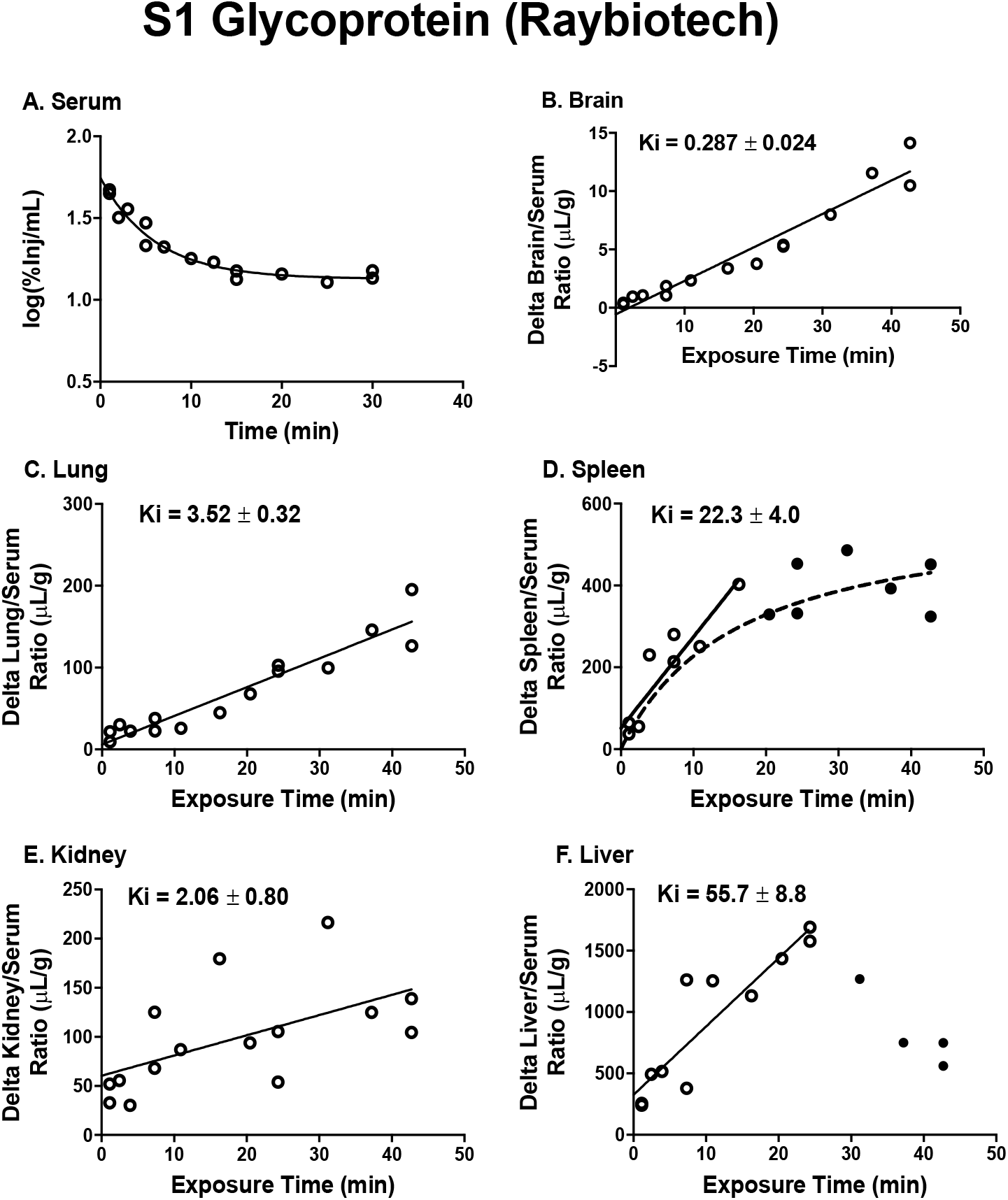
Uptake by brain and peripheral tissues and clearance from blood of I-S1 protein (Raybiotech). Panel A shows clearance from blood fitted to a one phase decay model. Panel B shows brain/serum ratios corrected for T-Alb (delta brain/serum ratios) plotted against exposure time according to multiple-time regression analysis. The slope of the linear portion of this relation measures Ki, the unidirectional influx rate. Panels C-F show the unidirectional influx rates for peripheral tissues (see Results for statistics).

Table 1 illustrates the stability of I-S1 from Raybiotech in blood and brain as measured by acid precipitation. Processing controls show the methodology did not itself result in degradation. Some degradation did occur in blood with an acid precipitation value (uncorrected by the processing control) of about 72%. In contrast, brain showed good acid precipitation (~77%) at 10 min but was at about 24% by 30 min. This likely represents, at least in part deiodinase activity, which is very concentrated in the brain, as well as degradation of the S1 protein itself in brain. Nevertheless, the results indicate that brain kinetics are best calculated based on data taken within a short time after iv administration.

**Table 1.** Stability as Assessed by Acid Precipitation of I-S1 in Brain and Blood 10 and 30 min after Intravenous Injection.

|  | 10 min | 30 min | Processing Control |
| --- | --- | --- | --- |
| Blood | $98 \pm 0.8$ | $72 \pm 1.8$ | $94 \pm 0.3$ |
| Brain | $77 \pm 0.04$ | $24 \pm 5.5$ | $95 \pm 0.2$ |

Values are ± SE; n = 3 for 10 min and 30 min and n = 2 for Processing Control.

Figure 3 shows the clearance of I-S1(Amsbio) from blood and its uptake by brain and other tissues. In general, patterns were very similar to the I-S1 from Raybiotech, but there were a few differences. The half-time clearance rate during the first 10 min after injection was 3.6 min. Data for the entire 30 min also was fit to a one phase decay model: Y0 = 1.98, plateau = 0.880, K = 0.393, and half life = 1.74 min, r^2^ = 0.867. Thus, clearance was somewhat faster for Amsbio I-S1 and the volume of distribution in the body somewhat less. Brain had a Ki = 0.294 ± 0.032 μL/g-min, r^2^ = 0.902, p<0.0001 (Figure 3B). Uptake patterns and rates are shown in Figure 3 for lung (3C, r^2^ = 0.896, p<0.0001), spleen (3D, r^2^ = 0.808, p = 0.002), kidney (3E, r^2^ = 0.765, p = 0.0004), and liver (3F, r^2^ = 0.883, p = 0.0005). Only liver showed a nonlinear pattern of uptake and uptake was nearly 5 times greater in liver for Amsbio I-S1 compared to Raybiotech S1. Some of the delta values for Amsbio I-S1 were negative at the early time points, indicating that the S1 brain/serum ratio was lower than the T-Alb brain/serum ratio at those time points. For Amsbio I-S1, values for lung, spleen, and kidney were very similar, whereas I-S1 from Raybiotech showed values for spleen that were about 10 times greater than that of lung and kidney.

**Figure 3.**
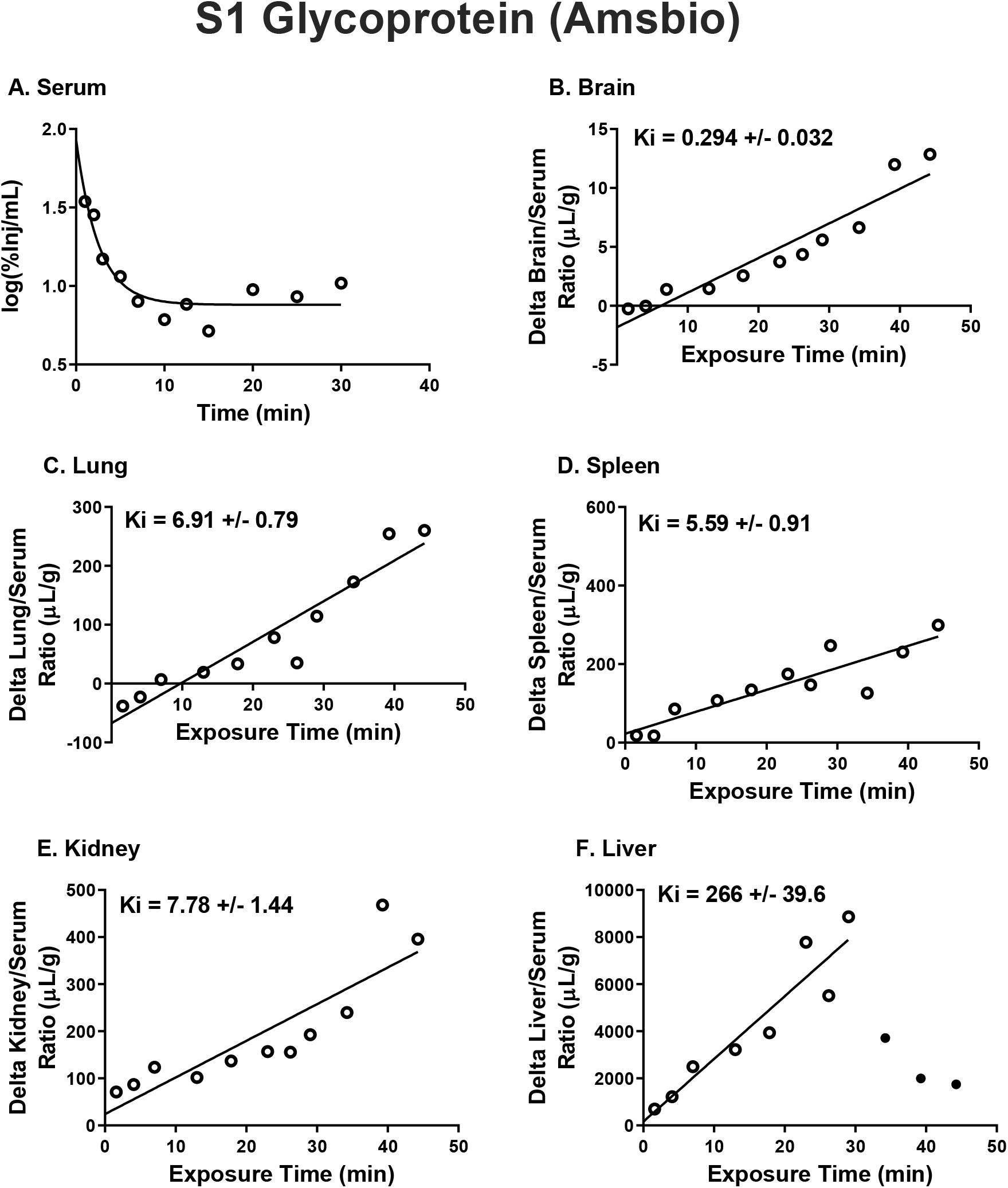
Uptake by brain and peripheral tissues and clearance from blood of I-S1 protein (Amsbio). Panel A shows clearance from blood fitted to a one phase decay model. Panel B shows brain/serum ratios corrected for T-Alb (delta brain/serum ratios) plotted against exposure time according to multiple-time regression analysis. The slope of the linear portion of this relation measures Ki, the unidirectional influx rate. Panels C-F show the unidirectional influx rates for peripheral tissues (see Results for statistics).

Capillary depletion is a quality control method to assure that material taken up by the BBB is completely crossing the BBB rather than being sequestered by the capillary bed. We used a modified technique of this method that also allows us to measure the amount of material that is reversibly adhering to the luminal side of the capillary bed. Figure 4 shows that the amount of I-S1 (Raybiotech) increased with time in the parenchymal (m = 0.187 ± 0.016 μL/g- min, r^2^ = 0.952, n = 9, p<0.0001) and capillary (m = 0.027 ± 0.006 μL/g-min, r^2^ = 0.757, n = 9, p = 0.0023) fractions, whereas it remained constant for luminal binding (r^2^ = 0.145) with a mean of 1.75 ± 0.50 μL/g.

**Figure 4.**
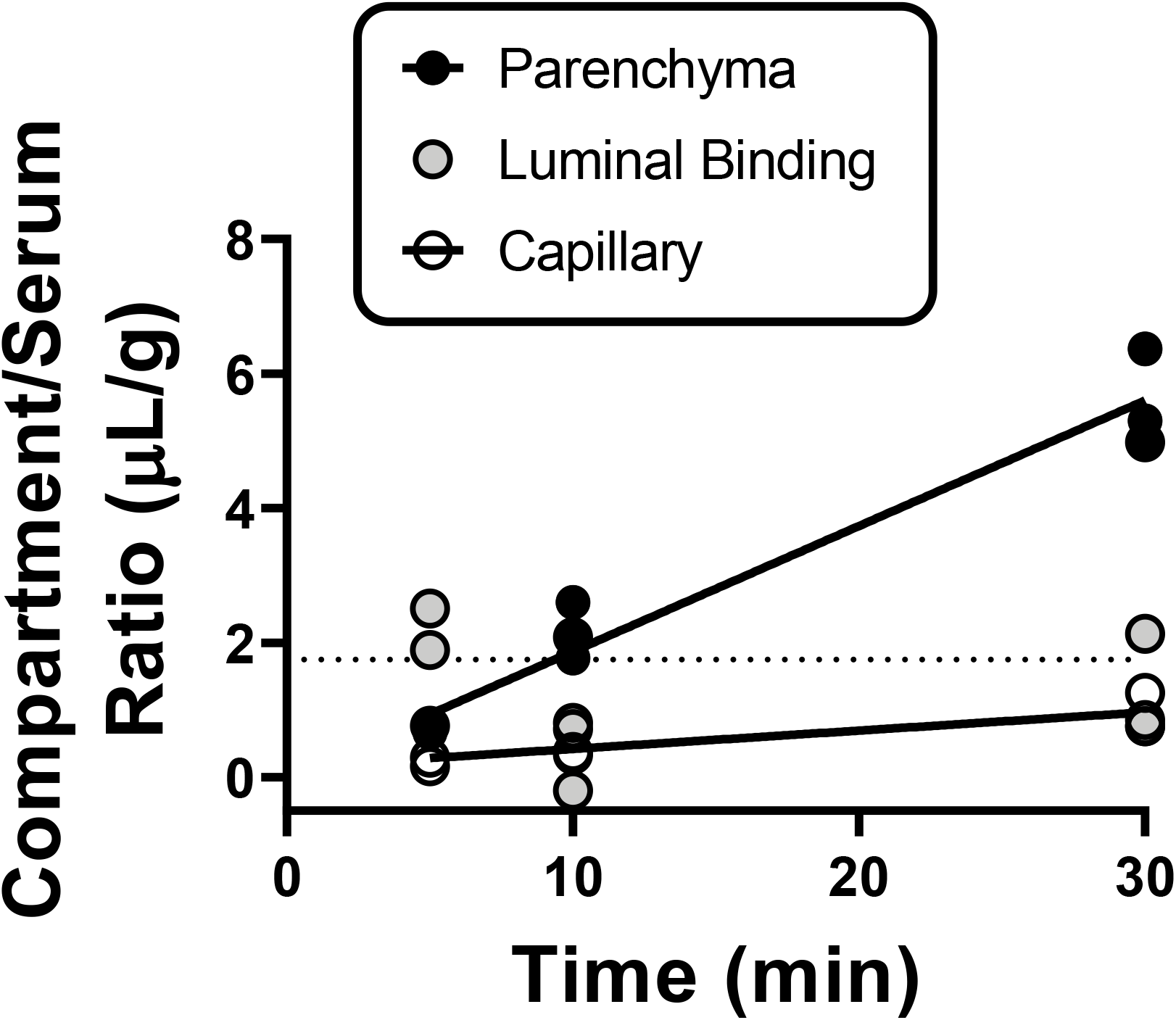
Capillary depletion for I-S1 protein. This method showed that I-S1 completely crossed the BBB to enter the parenchymal space of the brain and that this value increased with time. I-S1 was also detected in the capillary bed (blood-brain barrier) and also slightly increased with time, whereas luminal binding was steady at 1.75 μL/g (dashed line) during the course of the experiment.

Wheatgerm agglutinin (WGA) is a plant lectin that facilitates uptake of many glycoproteins, including those of viruses, and crosses the BBB by the mechanism of adsorptive transcytosis ^18, 37, 38^Here, we used 2-way ANOVAs followed by multiple comparisons test to compare effects on WGA treatment on the two sources of S1 proteins (n = 7 for each vehicle-treated group, n = 8 for WGA-treated Raybiotech group, n = 6 for WGA-treated Amsbio group). Figure 5 shows the effect of WGA on uptake by brain and peripheral tissues for I-S1 from both Raybiotech and Amsbio. Figure 5A also shows serum values and the effect of WGA on I-S1 from Raybiotech suggests increased clearance from blood compared to I-S1 from Amsbio (S1 source, treatment, interaction all significant at p<0.0001; multiple comparisons showed Raybiotech I-S1 different from all other groups at p<0.0001 with no other groups being different). For brain (Figure 5B), S1 source, treatment, and interaction were all significant at p<0.0001. WGA increased brain uptake for both I-S1’s (Raybiotech: p = 0.0007; Amsbio: p<0.0001), with a greater effect on Amsbio S1 (p<0.0001). Two-way ANOVA for lung (Figure 5C) showed an effect for WGA-treatment (p<0.0001), and multiple comparison’s test showed an increase for each I-S1 with WGA treatment (both at p<0.0001), but no difference between the two I-S1’s. For spleen (Figure 5D), two-way ANOVA showed an effect for S1 source and treatment, both at p<0.0001 and a trend for interaction (p = 0.07). WGA increased uptake for Raybiotech I-S1 (p<0.0001) but only induced a trend for Amsbio I-S1 (p = 0.08); the difference between the two WGA-treated I-S1 groups was significant (p<0.0001). For kidney (Figure 5E), two-way ANOVA showed differences for S1 source and treatment, both at p<0.0001. WGA increased uptake by kidney of both Raybiotech I-S1 (p = 0.0002) and Amsbio I-S1 (p = 0.003) with a difference between the two WGA-treated groups (p = 0.0005). For liver (Figure 5F), two-way ANOVA showed a significant effect for source of S1, WGA treatment and interaction, all at p<0.0001. Multiple comparisons test showed WGA significantly decreased uptake of Amsbio I-S1 (p<0.0001) but not Raybiotech I-S1 with no difference between the two WGA-treated groups.

**Figure 5.**
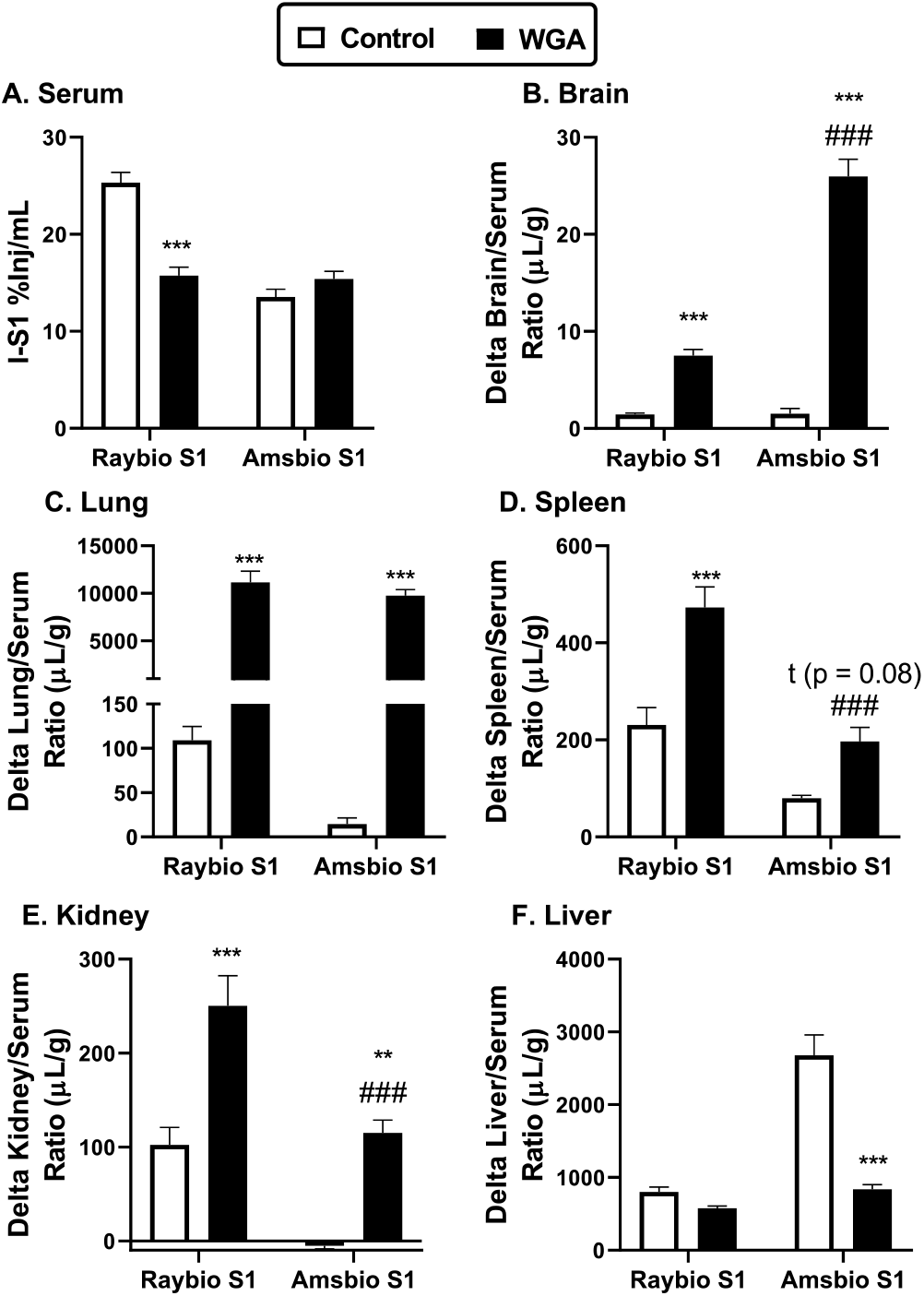
Effects of wheatgerm agglutinin (WGA) on I-S1 protein uptake. Values have been corrected for albumin space using T-Alb that was co-injected with I-S1. *, **, *** = difference between WGA vs no WGA for the indicated S1 at the p<0.05, <0.01, or <0.001 level; #, ##, ### = difference between Raybiotech vs S1 from Amsbio for WGA at the p<0.05, <0.01, or <0.001 level. For spleen, there was a trend (p = 0.08) for a difference between WGA and no WGA for the Amsbio I-S1.

It was next determined whether S1 transport into the brain could be saturated by excess unlabeled S1 protein, or if transport could be affected by ACE2 or its substrates AngII or ghrelin. The effects of 1ug/mouse of unlabeled proteins were compared by one-way ANOVA followed by multiple comparisons test with comparison to the vehicle groups only. None of these substances affected the level of delta I-S1 or T-Alb in blood, indicating that these substances did not affect the volume of distribution or clearance of either S1 or albumin. ACE2 did increase the renal level of T-Alb: F(4,36) = 2.63, p = 0.0505; vehicle: 119 ± 4.4 μL/g (n = 8); ACE2: 151 ± 15.6 μL/g (n = 8), p = 0.02; (data not shown). ACE2 also increased uptake of delta I-S1 by whole brain: F(4, 36) = 6.45, p = 0.0005; vehicle: 3.19 ± 0.24 μL/g (n = 8); ACE2: 5.24 ± 0.85 μL/g (n =8), p = 0.008 (Figure 6A). Additionally, S1 by t-test decreased uptake of I-S1 by whole brain (p = 0.026, f = 2.48, df = 14). Lung uptake of delta I-S1 was affected by all compounds except AngII (Figure 6B; note: one outlier of 201 μl/g was removed from the ghrelin group): F(4, 35) = 5.12, p = 0.0023; vehicle: 31 + 4.2 μL/g (n = 8); ACE2: 66 + 8.1 μL/g (n =8), p = 0.0007; S1: 53 + 5.5 μL/g (n =8), p = 0.037; ghrelin: 56 + 7.4 μL/g (n =7), p = 0.019. None of the substances affected the uptake of I-S1 by liver, kidney, or spleen (data not shown).

**Figure 6.**
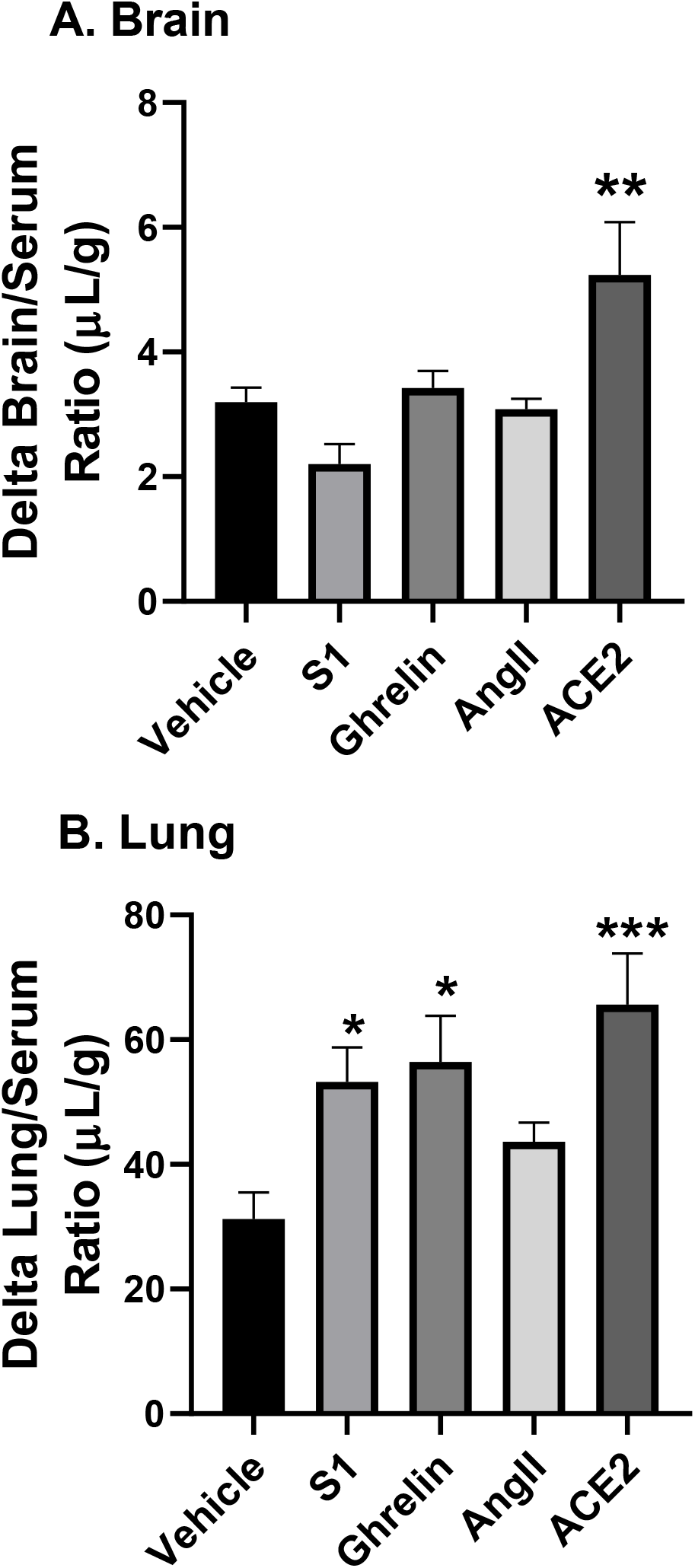
Competition and modulation of I-S1 protein uptake. Panel A shows that co-injection of unlabeled human ACE2 enhanced transport of I-S1 across the BBB. Panel B shows that I-S1 uptake by lung was enhanced by unlabeled S1, ghrelin, and ACE2. Uptake by olfactory bulb, liver, spleen, and kidney were not affected by any of these agents (data not shown).

Figure 7 shows uptake of I-S1 (Raybiotech) by brain regions; these values are the vehicle controls for the inflammation study (see Figure 8). One-way ANOVA showed a trend (p = 0.09) and multiple comparisons tests comparing all regions to one another or comparing to whole brain did not show any statistically significant differences or trends among the regions.

**Figure 7.**
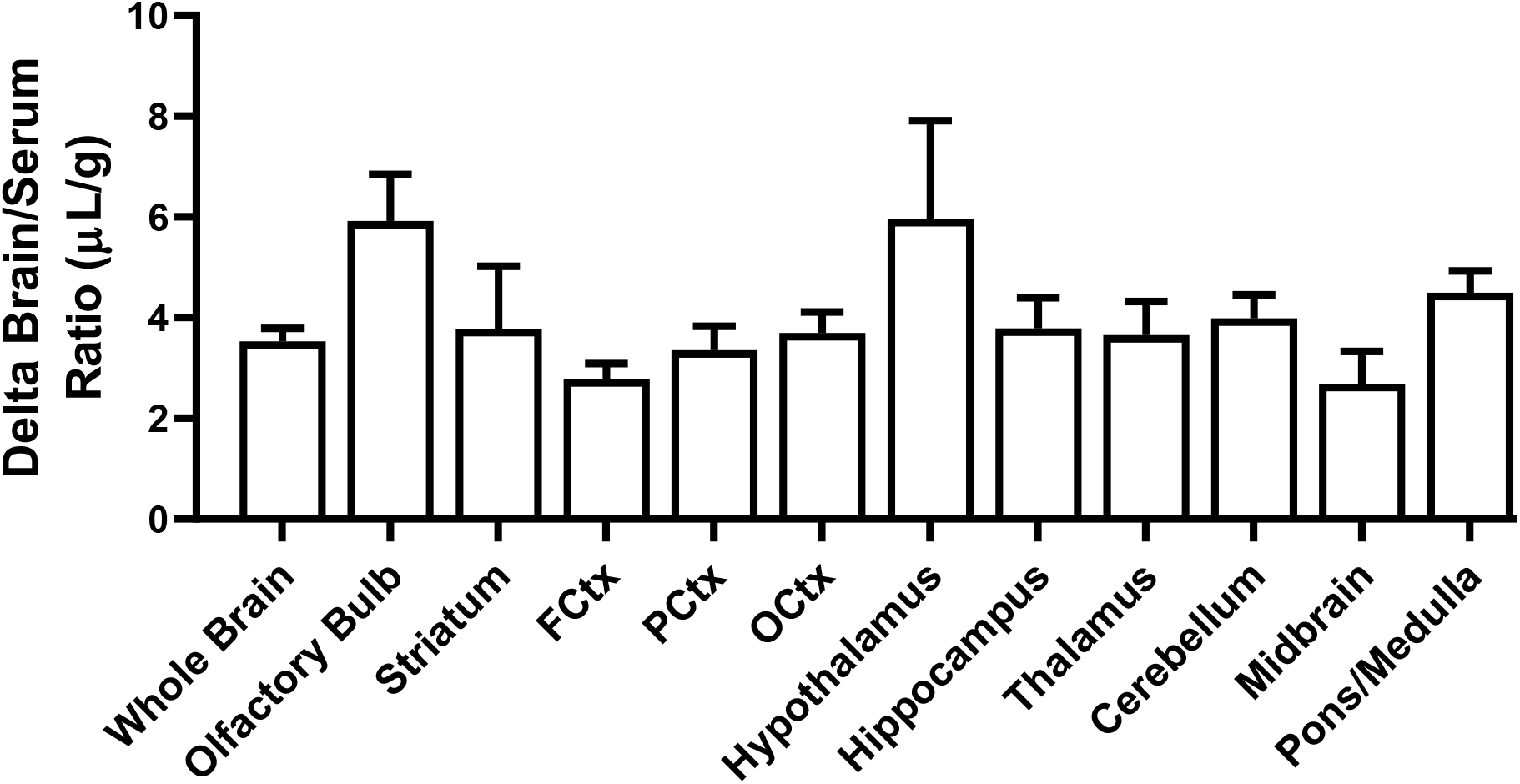
Brain regional uptake of I-S1 protein across the BBB. Data is from the controls from the Inflammation experiment in figure 8. ANOVA showed no statistically significant difference among brain regions.

**Figure 8.**
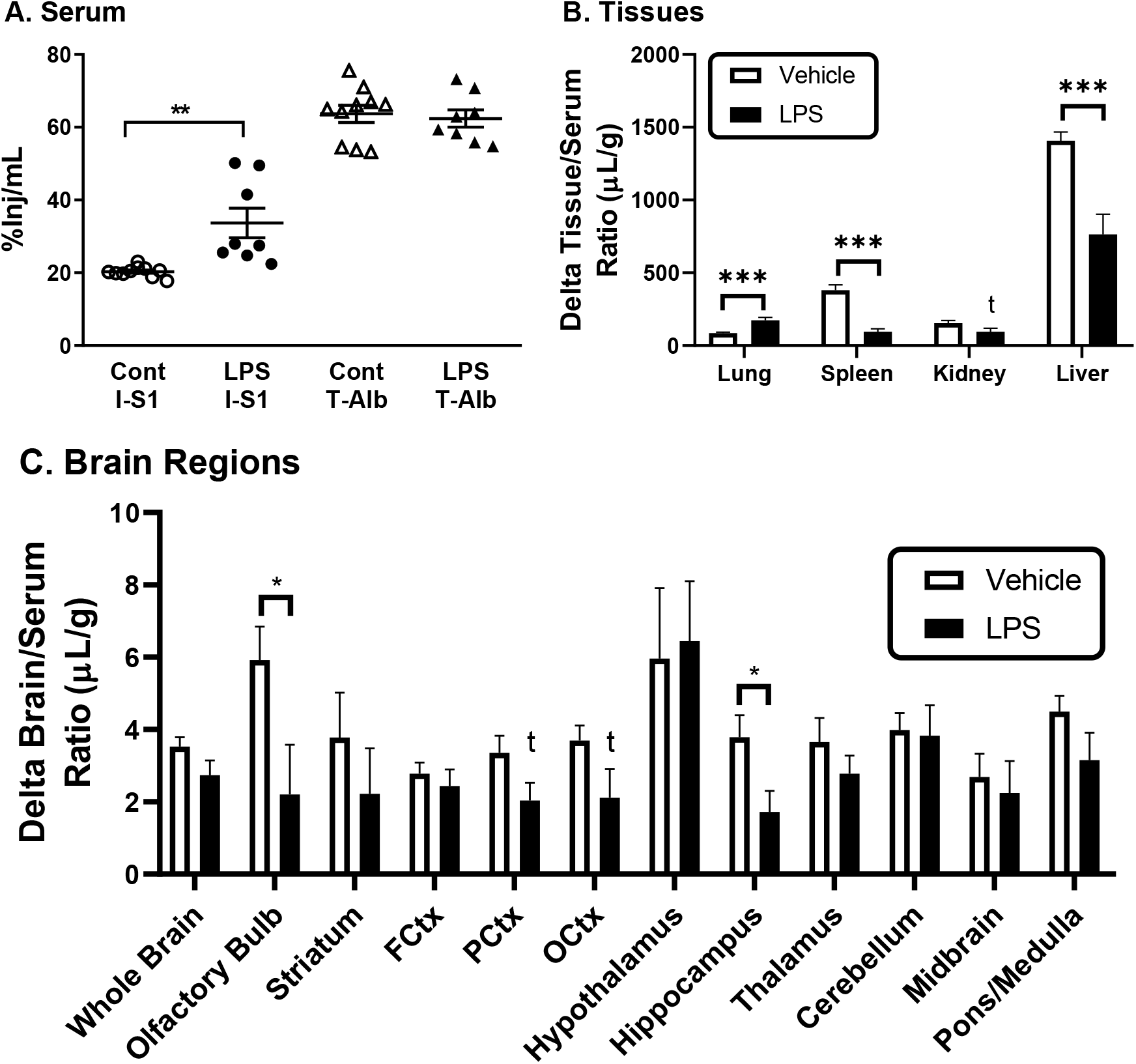
Effect of inflammation induced by LPS on I-S1 uptake. Panel A shows that treatment with LPS increased level of I-S1 in blood, indicating a decrease in the volume of distribution and/or clearance, but with no effect on T-Alb levels. Panel B shows that LPS treatment resulted in increased uptake of I-S1 by lung and decreases in uptake by spleen and liver, with a trend towards a decrease by kidney. Panel C shows levels in brain regions were arithmetically decreased, but reached statistical significance only in olfactory bulb and hippocampus with at trend for the occipital cortex. In all panels, *, **, *** = p<0.05, <0.01, or <0.001, and t = 0.1>p>0.05, respectively.

Results of treating mice with LPS are shown in figure 8. Figure 8A shows that LPS treatment increased the %Inj/mL for Raybiotech I-S1 (t = 3.66, df = 16, p = 0.0021), indicating a reduction in volume of distribution or clearance. There was no effect of LPS treatment on T-Alb (p = 0.7), demonstrating that the LPS effect on I-S1 was not caused by changes in vascular space or leakage. LPS treatment significantly increased lung uptake of I-S1 (t = 4.58, df = 16, p = 0.0003), while decreasing uptake for spleen (t = 6.39, df = 16, p<0.0001) and liver (t = 4.60, df = 16, p = 0.0003) with a trend for decreasing it for kidney (p = 0.056) (Figure 8B). Two-way ANOVA showed a brain region effect (p = 0.0028) accounting for 12.5% of the variability, and for LPS treatment (p = 0.0012) accounting for 4.5% of the variability (Figure 8C). Multiple comparisons test that compared each vehicle-treated group to its LPS-treated group found a significant decrease with LPS treatment for olfactory bulb (p = 0.026). Comparing vehicle- and LPS-treated groups by t-test showed decreases with LPS treatment for olfactory bulb (p = 0.034) and hippocampus (p = 0.028) with a trend for parietal cortex (p = 0.08).

Radioactivity appeared in blood after the intranasal administration of I-S1, indicating that some of the material entered the blood stream (data not shown). The AUC for blood after intranasal administration was 9.42 (%Inj/ml)-min, compared to the AUC for blood after iv injection was 1430 (%Inj/ml)-min, giving a bioavailability for the nasal route of 0.66%. Radioactivity was found in all brain regions at both 10 min and 30 min after the intranasal administration of I-S1. A t-test showed significant increases between 10 and 30 minutes for whole brain, frontal cortex, cerebellum, midbrain, and pons, but not the other regions. ANOVA for 10 min values showed a significant effect F(11, 48) = 3.77, p = 0.0006, with olfactory bulb (p = 0.031) and hypothalamus (p = 0.0041) being different from whole brain (Figure 9A). ANOVA at 30 minutes was also significant F(11,48) = 4.33, p = 0.0002, but only the olfactory bulb (p = 0.0023) was different from whole brain (Figure 9B). Figure 9A also shows for comparison the value for whole brain expressed as %Inj/g after iv injection.

**Figure 9.**
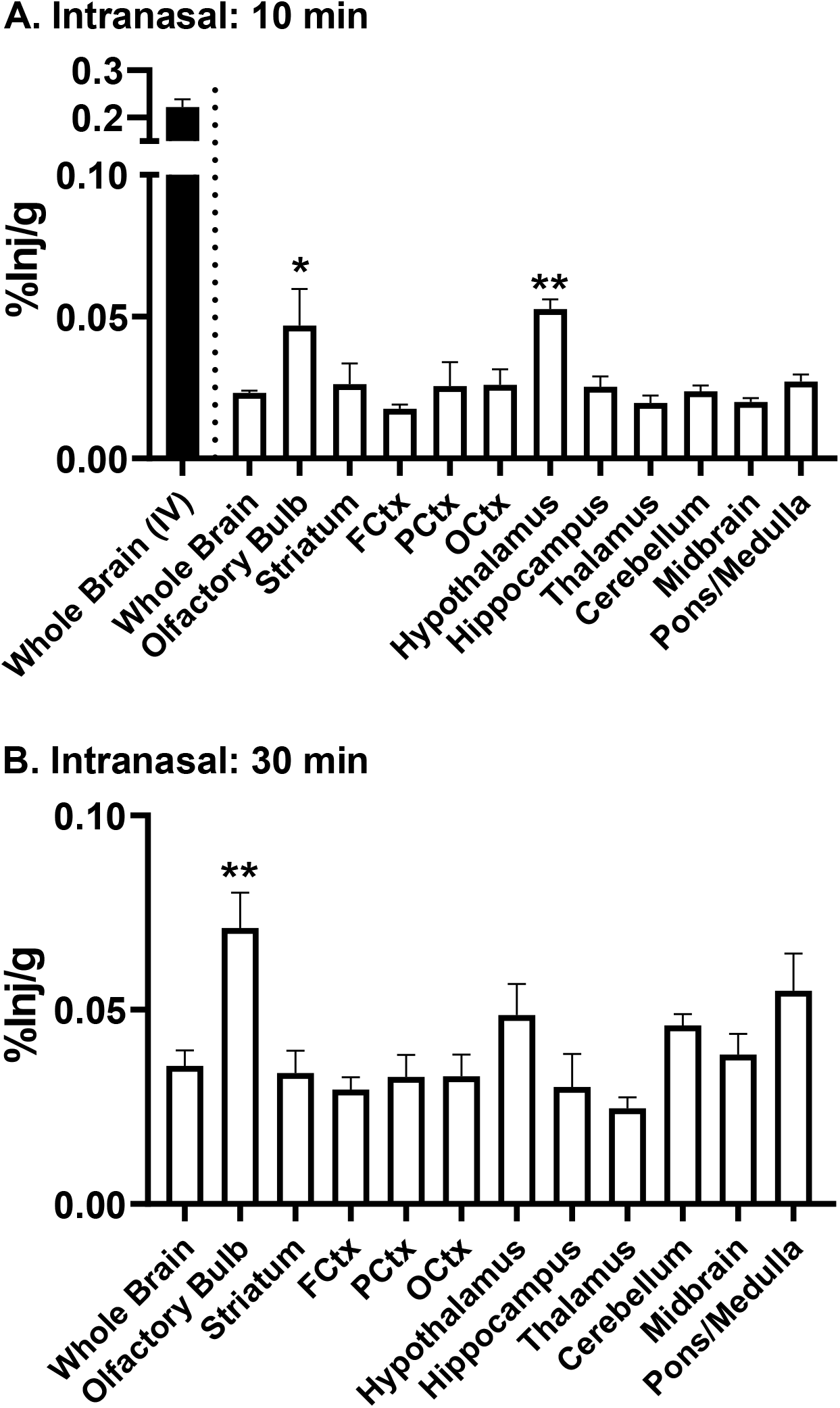
Brain regional uptake of I-S1 protein following intranasal administration. Panel A shows levels of I-S1 expressed as %Inj/g 10 min after intranasal administration and Panel B shows %Inj/g 30 min after administration for whole brain and brain regions. For comparison, Panel A also shows results for whole brain expressed at %Inj/g after intravenous injection of I-S1 (solid bar). In each panel, *, ** = p<0.05, <0.01, respectively.

The BBB was not disrupted in male or female ApoE3 or ApoE4 mice as evidenced by T-Alb spaces ranging from a mean of 7.7 to 8.4 μL/g in whole brain (Figure 10, Panel A). Clearance of I-S1 from blood (Figure 10, Panel B) and transport into whole brain (Figure 10, Panel C) and uptake by lung (Figure 10, Panel E) also did not vary as a function of ApoE genotype or sex. Comparison of slopes and intercepts with the Prism program showed a difference in slopes for the olfactory bulb, demonstrating that the unidirectional influx constant for I-S1 differed (Figure 10, Panel D): F(3,29) = 8.60, p = 0.0003. A two-way ANOVA of slopes showed a significant effect of sex (p = 0.0007) accounting for 29% of the variability; ApoE showed a trend (p = 0.06). Multiple comparisons for the olfactory bulb showed a significantly larger Ki for ApoE3 males vs ApoE3 females (p = 0.0056) and ApoE3 males vs ApoE4 females (p = 0.0015). For liver (Figure 10, Panel F), a significant difference occurred among the slopes: F(3, 20) = 5.06, p = 0.009). Two-way ANOVA showed a significant effect for ApoE (p<0.0001) accounting for 50% of the variability, and multiple comparisons showed a significantly lower uptake rate for ApoE4 females vs ApoE3 females (p = 0.0057), ApoE4 females vs ApoE3 males (p = 0.0001), and ApoE4 males vs ApoE3 males (p = 0.0039). For spleen (Figure 10, Panel G), a significant difference occurred among slopes: F(3, 24) = 17.4, p <0.0001. Two-way ANOVA showed a significant effect occurred for sex (p = 0.0004) accounting for 20% of the variability, for ApoE (p<0.0001) which accounted for 43% of variability, and for interaction (p= 0.0029) which accounted for 13% of variability, and multiple comparisons showed faster uptake rates for ApoE3 females vs ApoE4 females (p<0.0001), ApoE3 females vs ApoE3 males (p = 0.0003), and ApoE3 females vs ApoE4 males (p<0.0001). For kidney (Figure 10, panel H), comparison of slopes showed a difference: F(3, 22) = 4.51, p = 0.013. Two-way ANOVA showed a difference for ApoE (p = 0.0046) accounting for 25% of variability, and multiple comparisons showed faster uptake by ApoE3 males vs ApoE4 males (p = 0.0089) and ApoE3 males vs ApoE4 females (p = 0.012).

**Figure 10.**
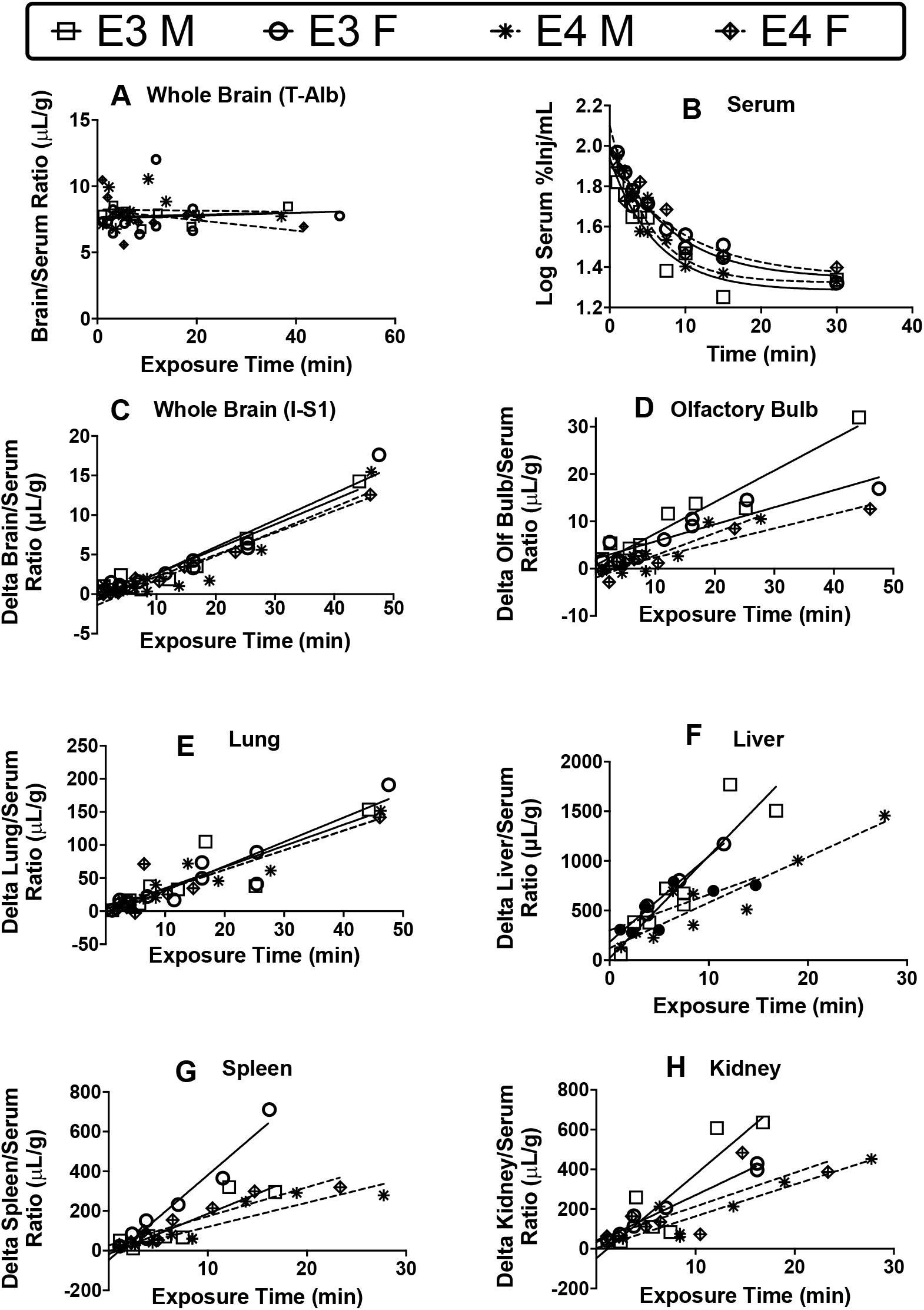
Effects of Sex and APOE genotype on clearance and uptake of I-S1 (Raybiotech). BBB integrity as measured by T-Alb space (Panel A), clearance of I-S1 from blood (Panel B), and uptake by whole brain (Panel C) and lung (Panel E) were not statistically affected by sex or ApoE status. Olfactory bulb was affected significantly by sex (Panel D). Liver (Panel F) and kidney (Panel H) affected by ApoE status. Spleen (Panel G) affected by ApoE status, sex, and interaction.

To determine whether the S1 transport system was intact in a human *in vitro* model of the BBB, we compared transport of I-S1 vs T-Alb across monolayers of iBECs seeded on transwells. For all studies, TEER values exceeded 1000 Ω*cm^2^ and TEER means were confirmed to be equal among groups just prior to starting the transport study. Three independent experiments were conducted with Raybiotech I-S1 to compare transport in the luminal to abluminal direction vs. T-Alb (Figure 11, panel A). Single experimental replicates evaluated Raybiotech I-S1 transport saturability with excess unlabeled S1, which was found to have no effect on transport of either I-S1 or T-Alb (Figure 11, panel B). WGA also had no effect on Raybiotech S1 transport (Figure 11, panel C). Raybiotech I-S1 had a significantly higher permeability coefficient when compared with Amsbio I-S1 *in vitro* (Figure 11, panel D), however this difference was not significant after correcting for T-Alb transport.

**Figure 11.**
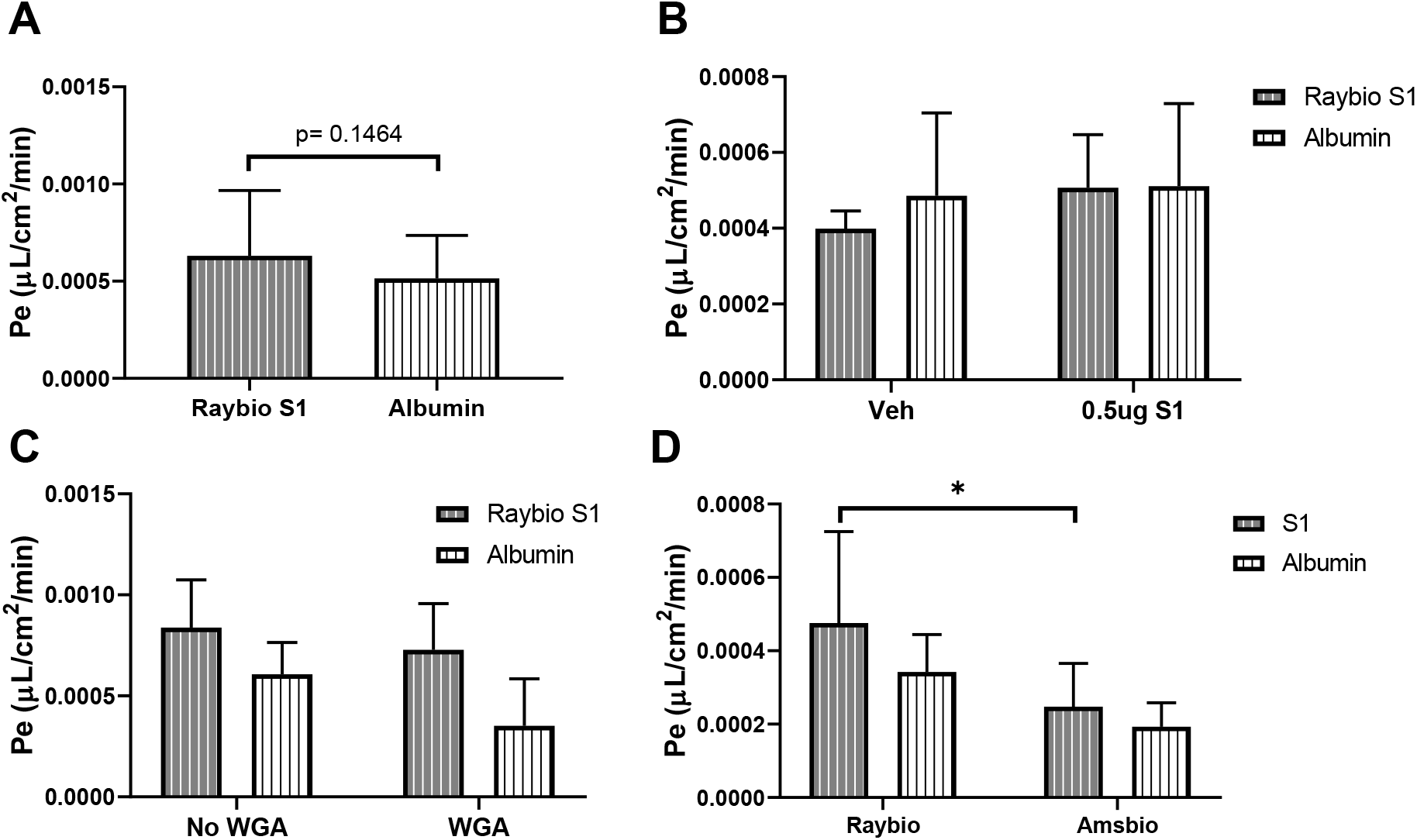
Transport of S1 proteins across human iBEC monolayers. I-S1 (Raybiotech or Amsbio) and T-Alb transport in the luminal to abluminal direction were studied. Panel A: Raybiotech I-S1 vs. T-Alb transport summarized from three independent differentiations, n=17 wells/group, average TEER (2578 ± 897.8 Ω*cm^2^). Panel B: Saturability of Raybiotech I-S1 transport with 0.5μg/well excess unlabeled S1, n= 5-6 wells/group. Panel C: Effects of WGA (0.5μg/mL) on Raybiotech I-S1 transport, n= 5-6 wells/group. Panel D: Comparison of Raybiotech and Amsbio I-S1 transport, n=6 wells/group, *p < 0.05.

## Discussion

These results clearly show that I-S1 from two different commercial sources readily and robustly cross the BBB. Other major findings of these studies were that uptake was rather uniform throughout brain, entering every region examined. I-S1 was also readily taken up by spleen, lung, and kidney with the highest uptake being by liver, indicating that to be the chief organ of clearance. The S1 protein completely crossed the BBB to enter the brain parenchymal space, with smaller amounts internalized by the brain endothelial cells and binding to their luminal surfaces. WGA greatly increased brain uptake of I-S1, indicating that S1 binds to sialic acid and/or N-acetylglucosamine and likely crosses the BBB by the mechanism of adsorptive transcytosis. Inhibition studies support adsorptive transcytosis and ACE2 as a binding site on brain endothelial cells. Our studies indicated ACE2 to also be important for lung, but not for liver, kidney, or spleen uptake. Inflammation tended to decrease clearance of I-S1 from blood, uptake by whole brain and brain regions, and the peripheral tissues except for lung, whose uptake was increased by inflammation. ApoE isoform and sex had no effect on uptake by whole brain, but males tended to transport I-S1 into the olfactory bulb faster than females. ApoE3 was associated with a higher uptake of I-S1 for liver, spleen, and kidney. Intranasally administered I-S1 entered both the brain and blood, but at much lower levels than after intravenous injection and the kinetics pattern indicates that the I-S1 in brain directly crossed the cribriform plate as opposed to entering from blood. Finally, in vitro studies showed similar, but attenuated, patterns to those found in vivo, suggesting that in vitro models of the BBB may be too dedifferentiated to be used in exploratory studies of S1 and SARS-CoV-2.

To improve specificity and statistical robustness of these studies, T-Alb was used as a control, being co-injected into every animal, thus measuring vascular space for brain and vascular space plus leakage for peripheral tissues in each individual, and allowing for delta tissue/serum levels to be calculated. These delta values, therefore, measure the uptake that is specific to I-S1. The transport rates, or Ki’s, as measured by the delta brain/serum ratios differed by less than 3%, with Ki = 0.287 + 0.024 μl/g-min for the S1 from Raybiotech and a Ki = 0.294 + 0.032 μl/g-min for the S1 from Amsbio. Likewise, the two I-S1’s had similar patterns for clearance from blood and uptake by peripheral tissues, with liver being by far the highest for both I-S1’s and having an early nonlinear component to uptake.

There were subtle differences between the kinetics of the two I-S1’s which are likely caused by differences in glycosylation since this is critical to the tissue uptakes of viral proteins ^39, 40^. The S1 from Raybiotech was cleared less rapidly from blood and had an early non-linear component to uptake by spleen and liver. The nonlinearity is usually caused by an efflux component from tissue back into the blood stream. This suggests that liver and spleen may be re-releasing previously sequestered I-S1 back into the blood stream. The I-S1 from Raybiotech showed more variation among uptake rates for lung, spleen, and kidney, and had a lower uptake by liver in comparison to the I-S1 from Amsbio. One consistent finding was that the Amsbio I-S1 often produced negative values for tissue/serum ratios, especially at early time points and for brain. This literally means that the vascular space for I-S1 was less than that for T-Alb at those time points. Given the larger size of S1 in comparison to albumin, it is not surprising that albumin would be more leaky at the peripheral tissues. However, it is less clear how negative values occurred for brain uptake.

Capillary depletion was used to follow the temporal pattern of uptake of I-S1 from Raybiotech into the brain space. Occasionally, substances accumulate either on the luminal surface of the brain or are sequestered by the brain capillaries without subsequent passage into brain tissue. If those processes occur over time, then multiple-time regression analysis can give the false impression that penetration of the BBB is occurring. Here, capillary depletion showed a strong time-dependent uptake of I-S1 into brain tissue and a much slower increase in the amount of I-S1 retained by brain capillaries. In contrast, the amount of I-S1 reversibly binding to the luminal surface of brain endothelial cells was constant. This shows that luminal binding to brain endothelial cells quickly reached a steady state with blood levels, and that most of the I-S1 taken up by brain endothelial cells quickly entered the brain parenchymal space. Thus, brain cells are exposed to S1 which could then bind to or activate them. Similarly, S1 internalization could affect brain endothelial cell function and provides a mechanism by which S1 could facilitate infection of brain endothelial cells.

WGA is a glycoprotein plant lectin that binds to sialic acid and to N-acetylglucosamine, inducing adsorptive transcytosis in brain endothelial cells, a vesicular process that results in transport across the BBB ^41, 42^. Co-injection of WGA increased uptake by brain, lung, spleen, and kidney for I-S1 from either Raybiotech or Amsbio. WGA decreased serum values for I-S1 from Raybiotech, indicating increased volume of distribution among tissues or clearance from blood, and decreased liver uptake for I-S1 from Amsbio. The endosomes induced by adsorptive transcytosis are often routed to the lysosomal compartment and viruses which cross the BBB by the mechanism of adsorptive transcytosis, including HIV-1 and rabies, can survive the lysosomal compartment ^37, 43–45^. An interesting feature of adsorptive transcytosis is that when a robust stimulator such as WGA is co-injected with a less robust stimulator, WGA promotes, rather than inhibits, the transport of the less robust stimulating substance. For example, co-injection of WGA with HIV-1 gp120 greatly increases the transport of gp120 across the BBB ^46^. Thus, the finding that WGA increased uptake by brain is consistent with adsorptive transcytosis as a mechanism of entry. These results suggest that S1, and by extension possibly SARS-CoV-2, may cross the BBB or be taken up by peripheral tissues by binding to membrane bound glycoproteins that contain sialic acid and N-acetylglucosamine.

Subsequent studies used S1 from Raybiotech except as noted.

The major cell surface protein that S1 is thought to bind to is ACE2 ^47, 48^. Based on experience with SARS, it has been assumed that SARS-CoV-2 will bind to human, but not murine, ACE2. SARS can infect mice, but doesn’t produce severe symptoms and death except in transgenic mice overexpressing human ACE2. However, at least part of this effect in transgenic mice may be caused by their expressing 8-12 times more ACE2 ^49^. The spike protein of SARS and SARS-CoV-2 share 77% sequence identity with the S1 from SARS-CoV-2 having 4 more charged residues ^47^. These extra positive charges would facilitate transport across the BBB by adsorptive transcytosis ^50, 51^. Mouse and human ACE2 are 84.2% homologous ^52^. Therefore, it should not be prematurely assumed that the S1 from SARS-CoV-2 cannot effectively bind to mouse ACE2. However, as ACE2 expression is low in brain ^53, 54^, it may be that other receptors are used to enter brain. This would be analogous to HIV-1, which does not use its classic binding sites to cross the BBB ^55, 56^.

S1 binds off-target from the ACE2 catalytic site ^57^. Here we tested the ability of ACE2 to compete with brain uptake of I-S1. We co-injected with I-S1 either soluble ACE2, unlabeled S1, or the ACE2 substrates ghrelin or angiotensin II. Only ACE2 affected transport across the BBB and its presence stimulated uptake. One would have predicted that soluble ACE2 would have decreased I-S1 transport across the BBB by competing with the ACE2 present on brain endothelial cells. Short term, soluble receptors for a substance typically inhibit BBB transport, although they can ultimately increase total brain uptake ^58^. Lung produced more paradoxic findings in that unlabeled S1 and ghrelin as well as ACE2 produced statistically significant increases in uptake. These findings, although paradoxical, suggest that ACE2 is involved in uptake of I-S1 by both the BBB and lung and the paradoxical increases they induce are consistent with transport by the mechanism of adsorptive transcytosis.

AngII may have been ineffective in alteringI-S1 uptake by any tissue since I-S1 binds to a different location on ACE2 ^47^. This is typical of viral proteins which usually don’t bind to the active site of a transporter or enzyme, but target some less specific aspect of the sugar-protein complex. Viral proteins are less discriminating than the ligands of the glycoproteins they target and so may bind to a variety of membrane glycoproteins. This is exemplified by rabies and herpes simplex viruses each of which binds to multiple receptors ^59, 60^. The lack of effect of the tested substances on kidney, liver, and spleen may suggest that other receptors are involved in S1 sequestration. Receptors besides ACE2 can bind SARS-CoV-2, including basigin, cyclophilins, and dipeptidyl peptidase-4 ^61^ and GRP78 ^62^.

Uptake of I-S1 was very uniform throughout brain regions with no statistically significant differences among brain regions. The region with the highest uptake was the hypothalamus at 6.0 μl/g and the region with the lowest uptake was the midbrain at 2.6 μl/g. This widespread uptake throughout the brain supports the consideration that the central nervous system may be contributing to many of the diverse effects of S1 and/or SARS-CoV-2 such as encephalitis, respiratory difficulties, and anosmia ^1, 3, 4^.

Inflammation can increase the passage of substances across the BBB by a variety of mechanisms, including stimulation of adsorptive transcytosis. For example, the LPS treatment used here greatly increases the uptake of both gp120 and HIV-1 by brain ^63, 64^. LPS releases many of the same cytokines associated with the cytokine storm of COVID-19, including interleukin (IL)-6, IL-10, granulocyte colony-stimulating factor, granulocyte-macrophage colony-stimulating factor, interferon gamma-induced protein 10, monocyte chemoattractant protein-1, and tumor necrosis factor-alpha ^5, 65^. Here, LPS increased uptake of I-S1 by lung. This would suggest that the cytokine storm of COVID-19 could be associated with increased uptake of S1 and SARS-CoV-2 by lung. Otherwise, LPS tended to decrease, not increase, uptake by tissues. LPS increased blood levels of I-S1, consistent with a reduction of uptake by tissues or decreased clearance. Indeed, treatment with LPS reduced uptake by spleen and liver with a trend for kidney. Most brain regions had an arithmetic, non-statistically significant decrease in I-S1 uptake with LPS treatment, although two regions (olfactory bulb and hippocampus) did reach statistical significance. These results suggest that the cytokine storm produced by LPS may, at least as far as S1 actions are concerned, be beneficial for peripheral tissues such as liver and spleen, neutral to beneficial for brain, and harmful for lung.

Viruses present in the nasopharyngeal cavity can enter the brain. Viruses in the nasal passages can be absorbed directly into the blood stream, especially by the very vascular turbinates, using the hematogenous route to infect peripheral tissues and the brain ^40^. Other substances can enter the central nervous system by being absorbed at the cribriform plate and by nerve terminals of the trigeminal nerve ^66^. Here, we administered I-S1 at the level of the cribriform plate where it has been shown substances can be taken up by the cerebrospinal fluid, olfactory bulb, and subsequently other brain regions ^67^. We found I-S1 present in all brain regions at 10 min, with olfactory bulb and hypothalamus having the highest levels and brain levels even higher at 30 min. Radioactivity was also found in blood 10 and 30 min after the intranasal administration. Both brain and blood levels after intranasal administration are much lower than after intravenous administration, with brain levels after intranasal administration being 1/10^th^ and blood levels about 1/150^th^ of levels after intravenous administration. I-S1 could have first entered into the blood stream with subsequent blood-to-brain transmission, or have first entered the brain with subsequent entry into blood with the reabsorption of cerebrospinal fluid. However, the blood levels produced after intranasal administration are far too low to account for more than about 1/15^th^ of the radioactivity found in brain. Therefore, the results support I-S1 entry into the central nervous system across the cribriform plate or by the trigeminal nerves. However, the amount of radioactivity entering the brain was much greater after intravenous than after intranasal administration. This suggests that although S1 and SARS-CoV-2 can enter the brain directly from the nasal passages, the major route is likely hematogenous.

The ApoE4 genotype and being male has been associated with increased risk of severe COVID-19 symptoms ^20, 21^. Here, sex and ApoE genotype exerted no statistically significant effect on I-S1 clearance from blood or uptake by whole brain or lung. Sex did affect uptake of I-S1 for olfactory bulb, with males having a higher uptake, and for spleen, with males having a lower uptake. ApoE genotype affected uptake by spleen and kidney with a trend for olfactory bulb. In all cases where significant, ApoE3 was associated with a higher uptake of I-S1 than was ApoE4. ACE2 levels are lower in humans expressing ApoE4 and so would be expected to have lower levels of viral uptake ^68^. It should be noted that the ApoE studies were conducted in 4 mo old mice and most of the effects associated with ApoE4 occur with advancing age. Therefore, it is important to examine S1 and viral uptake in aged ApoE isoform mice.

In comparison to in vivo studies, the vitro studies did not produce robust results for I-S1 uptake. Transport of I-S1 was not observed to be significantly greater than that of albumin in a human in vitro model of the BBB, nor was it affected by unlabeled S1 protein or WGA. Therefore, the transport mechanisms that are intact for I-S1 in vivo in mice are not present or are attenuated for I-S1 in our iBEC model despite findings of other BBB-characteristic features such as high TEER and consistent expression of brain endothelial cell markers such as claudin-5, GLUT-1, and PECAM, similar to what has been shown by others ^35^. Our findings indicate that model optimization is required for studies of S1 transport in vitro using iBECs. They also show that the transport of I-S1 across the BBB found in vivo depends on more selective mechanisms such as glycoprotein binding rather than less selective such as leakage.

In summary, these studies clearly show that I-S1 glycoprotein crosses the BBB and is taken up by peripheral tissues. The tissue with the highest uptake by far is liver, suggesting it is the tissue that primarily clears S1. I-S1 completely crosses the BBB to enter brain tissue and enters in all regions of the brain. WGA agglutinin enhances uptake of I-S1 by brain and by peripheral tissues, suggesting transport across the BBB involves sialic acid or N-acetylglucosamine and likely uses the transcytotic mechanism of adsorptive transcytosis. ACE2 given intravenously with I-S1 paradoxically increases brain uptake of I-S1 as well as that of lung, but not for kidney, liver, or spleen, suggesting S1 may use other membrane glycoproteins in addition to ACE2 as functional receptors. Inflammation tends to reduce uptake of I-S1 by brain regions and for peripheral tissues except lung, suggesting a protective effect. Against S1. Sex and ApoE have no effect on I-S1 uptake by whole brain and lung, but do have tissue dependent effects for olfactory bulb, liver, spleen, and kidney, with ApoE3 status associated with increased uptake of I-S1. Taken together, the findings here demonstrate and characterize the uptake of S1 by brain and peripheral tissues.

## Acknowledgements

Supported by the VA (WAB, MAE), NIH R21 AG065928-01 (MAE), and NIH RF-1 AG059088 (WAB, JR).

We wish to thank Kristen Baumann, Riley Weaver, Sarah Pemberton, and Danny Quaranta for technical assistance.

Graphical abstract created with BioRender.Com

---

**Supplemental Figure 1.**
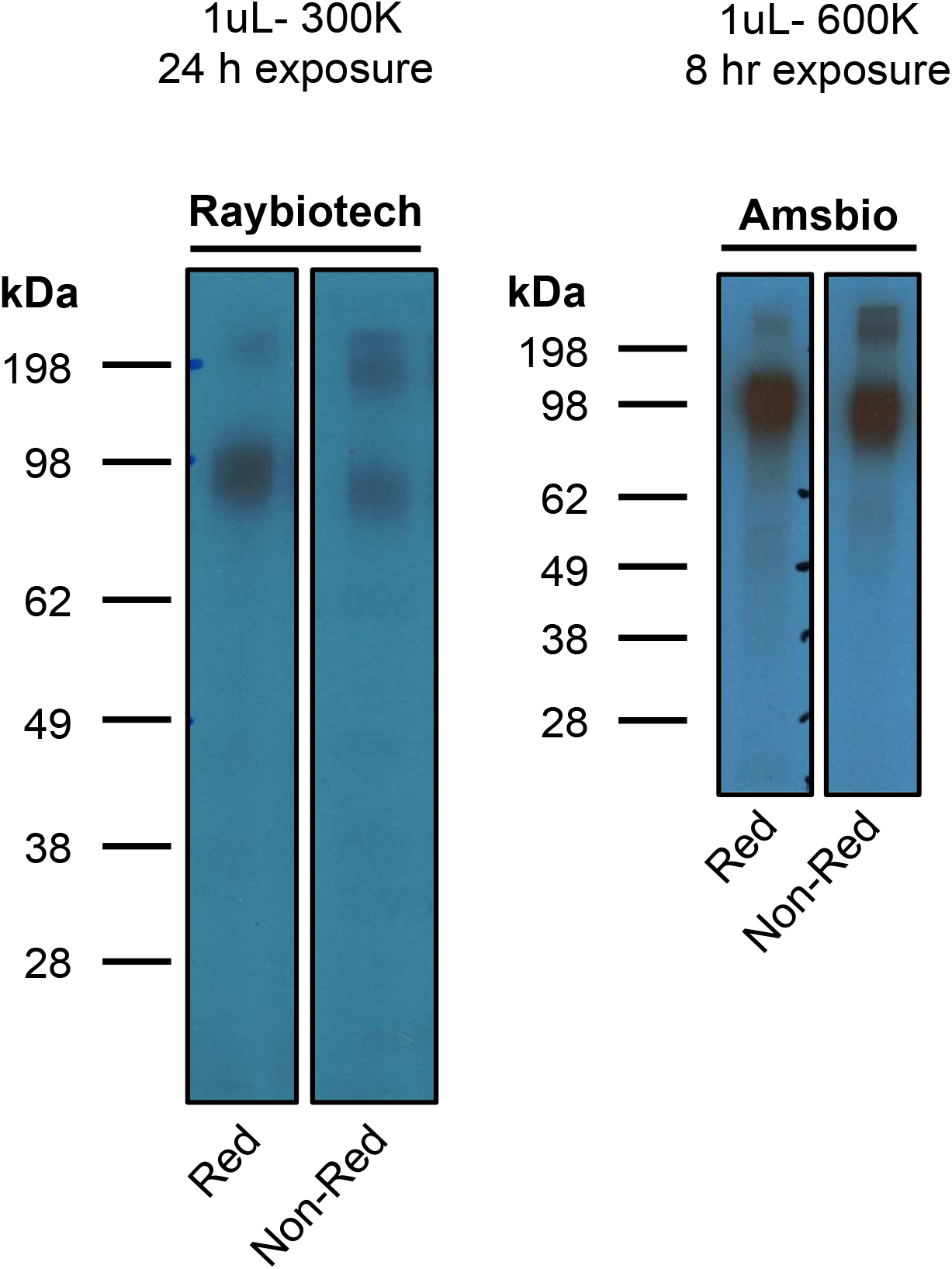
Gel Autoradiography of I-S1 from Raybiotech and Amsbio. In each case, both the reduced and non-reduced gels migrated at the molecular weights predicted by the manufacturers.

## Notes

### Competing Interest Statement

The authors have declared no competing interest.

